# HGNNPIP: A Hybrid Graph Neural Network framework for Protein-protein Interaction Prediction

**DOI:** 10.1101/2023.12.10.571021

**Authors:** Lutong Chi, Jinbiao Ma, Yingqiao Wan, Yang Deng, Yufeng Wu, Xiaochen Cen, Xiaobo Zhou, Xin Zhao, Yiming Wang, Zhiwei Ji

**Author notes:** Corresponding author: Zhiwei Ji, Yiming Wang. These authors contributed equally to this work (Lutong Chi, Jinbiao Ma).

## Abstract

A deep understanding of Protein-protein interactions (PPIs) can provide comprehensive insights into many biological functions, thereby facilitating drug target identification and novel therapeutic design. Recent developments in artificial intelligence (AI)-driven computational methods have enabled the discovery of previously uncharacterized PPIs from large-scale interactome datasets. Almost all existing machine learning methods rely on Subcellular Localization (SL) to construct balanced datasets based on positive interactions to achieve predictions. Despite high fitting accuracy, the generalization ability of these models is questionable. To solve this problem, we analyzed existing methods and found that the high false positives in these methods are due to the bias in data distribution caused by SL. Therefore, we proposed a new strategy for negative instance sampling in PPI prediction and developed a Hybrid Graph Neural Network framework for Protein-protein Interaction Prediction (HGNNPIP). The experimental results showed that HGNNPIP works well on six benchmark datasets. Comparison analysis demonstrated that our model outperformed the other four existing methods. We also used HGNNPIP to explore the molecular contacts involved in the rice-pathogen interaction system. *In vivo* experiments confirmed multiple regulations related to disease resistance in rice. In summary, this study provides new insights into establishing a computational framework for PPI prediction with high reliability.

## 1. Introduction

Systematic understanding of the protein-protein interactions (PPIs) can provide comprehensive insights into the molecular mechanisms of complex biological processes and diseases for different organisms ^1-3^. Discovery of new PPIs could facilitate important interventional goals, such as drug target identification and novel therapeutic design ^4^. Experimental technologies such as yeast two-hybrid ^5^, co-Immunoprecipitation (co-IP) ^6^, and fluorescence resonance energy transfer ^7^, *etc*. have provided information on the biophysical interactions between two or three proteins ^8,9^. However, the lack of coverage and quality of comprehensive PPI networks is one of the inherent shortcomings of the existing experimental approaches ^10^.

Computational approaches ^11-16^, especially those based on machine Learning (ML), can facilitate the identification of previously uncharacterized PPIs. ML-based algorithms can generally be divided into three categories: 1) supervised learning, 2) unsupervised learning, and 3) reinforcement learning. Among supervised learning methods, Bayesian inference ^17,18^, Decision tree ^19^, and Support Vector Machine (SVM) ^20^ are the most representative models for PPI prediction. Unsupervised learning strategies, including K-means ^21^ and Spectral clustering ^22,23^, are also widely used for PPI prediction. Unfortunately, the prediction performance provided by the above approaches appears to be unsatisfactory. Recently, reinforcement learning (RL) was used to detect new molecular complexes in PPI networks ^24,25^. The results from Palukuri, et al. showed that RL algorithm had good performance on the human PPI dataset ^24^. However, it is unclear whether it can be used in other scenarios.

In the past few years, many researchers have believed that deep learning (DL)-based methods can achieve better performance in PPI prediction compared with conventional machine learning-based models ^2,26-32^. Among these DL-based models, amino acid sequence, due to their easy accessibility, is the main data type of input. The representative amino acid sequence-based models include DeepFE-PPI ^28^, DPPI ^30^, PIPR ^29^, and DeepTiro ^27^. Until recently, topology information of PPI networks began to be integrated with protein sequences for PPI computational identification (e.g., S-VGAE ^31^; HNSPPI ^2^, *etc*.). To train the models with numerical data, various deep learning architectures were designed to implement feature embedding. Generally, Convolutional Neural Networks (CNN), Multilayer Perceptron (MLP), and Word2vec are widely used for sequence embedding ^27^. Graph learning methods, such as Graph Neural Network (GNN) and Graph attention network (GAN), have been applied to extract 3D structure features of proteins and network properties ^33,34^. Overall, the above well-known models appear to perform well on benchmark datasets (e.g., *Human* and *S. cerevisiae*) ^27-30,34^. However, their reliability in practical applications has rarely been reported. We believe that bias in data distribution and overfitting in model training may represent a severe limitation to the accurate identification of unknown PPIs.

In this paper, we first investigated three possible strategies used for negative sample generation in the PPI study and evaluated their impact on the prediction performance of trained models. All downloaded benchmark datasets were reconstructed before model training. We then developed a Hybrid Graph Neural Network framework for Protein-protein Interaction Prediction (HGNNPIP). HGNNPIP, as a hybrid supervised learning model, consists of sequence encoding and network embedding modules to comprehensively characterize the intrinsic relationship between two proteins. To verify the effectiveness of HGNNPIP, we tested it on the PPI datasets of six species. We also conducted a comparison analysis of the HGNNPIP model with the other four State-of-the-art (SOTA) algorithms. In addition, we used HGNNPIP to explore the molecular changes involved in the rice-pathogen interaction system and experimentally validated multiple regulations *in vivo*.

## 2. Methods

### Data collection and preparation

In this study, six PPI datasets were used for model evaluation and following analysis. These datasets were generated from multiple species, including *S. cerevisiae, C. elegans, E. coli, D. melanogaster, Human*, and *O. sativa*. The first four datasets were obtained from Guo’s work ^35^ and the original PPI data was extracted from a well-known database DIP (Database of interacting proteins) ^36^. The Human PPI dataset was collected from Pan’s work ^37^ and the corresponding data was referred to the database HPRD (Human Protein Reference Database) ^38^. In addition, we manually constructed a new PPI dataset for *O. sativa* based on STRING ^39^ (score>0.8), which was used in case study.

As shown in **Table 1**, the first five datasets contain 5594, 4030, 6954, 21975, and 36630 PPIs (positive instances), respectively. Both Guo et al. and Pan et al. demonstrated an equal number of negative instances based on protein subcellular localization information ^35,37,40^. Also, we used the same strategy to generate the negative set for the dataset *O. sativa*.

### The strategy for Negative Instance Generation

Although subcellular localization ^2^ is the most commonly used strategy to generate negative instances, it remains unknown whether the constructed negative set will cause obvious false positives. In this study, we proposed three potential strategies for negative instance generation and determined the best one for PPI prediction.

#### Popularity-biased Negative Sampling (PopNS)

As a heuristic strategy for negative sample distribution, Popularity-biased Negative Sampling (PopNS) was first proposed in the word2vec model ^41^ and is widely used in Recommendation systems (RS) ^42-44^. PopNS adopts a fixed distribution proportional to an item interaction ratio ^45^. In RS, Popularity is reflected by ‘frequency’ or ‘degree’ ^46^.

In this study, we defined the node degree in PPI network as the ‘popularity’ of a protein. Moreover, 2-hop and 3-hop neighbors are mainly focused as they are involved in local or long-distance relationships ^47^. Let’s illustrate how PopNS determines noninteracting pairs as negative instances from a given PPI network (**Supplementary Fig. 1**). The degree of a node is defined as the weighted sum of all connected links: 1 denotes a positive interaction (solid line with black color) and -1 denotes a negative interaction (dash line with red color).

For any node *v*, we first counted the interacting partners and obtained the initial degree. Then, the probability of each neighbor being a noninteracting partner was calculated. Negative Sampling is started by rolling a dice. Once a node is selected, a negative instance is added to the network, and the corresponding degree and probability are updated.

Here, we take *V*_1_ as an example. Before iteration, the search list contains five vertices: *V*_5_, *V*_6_, *V*_7_, *V*_8_, and *V*_9_. With the highest value *p, V*_7_ is selected as the first negative partner *V*_1_. As a negative edge is added to the network, the degree of *V*_1_ and *V*_7_ are modified as 2 and 3, respectively. Finally, the probability of the remaining vertices (*V*_5_, *V*_6_, *V*_7_, *V*_8_) are updated. Due to the higher value of *p, V*_6_ is then selected as the second negative partner *V*_1_. When the degree drops to 0, negative sampling for the vertex *V*_1_ is completed. Actually, our PopNS strategy assumes that if the current node is not a 1-hop neighbor of a node with a large degree, they are more likely to be determined as a negative pair.

#### Similarity-biased Negative Sampling (SimNS)

In recommendation systems, Similarity-biased negative sampling (SimNS) selects informative negative items based on the distances between positive and negative items through a two-stage strategy ^48-50^. This strategy takes into account the influence of positive items on negative sampling, which inspires us to incorporate the positive items into negative sampling ^50^.

In this study, we define the similarity between two proteins based on embedding vectors (equation (1)).

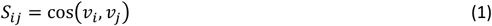

where *v*_*i*_ and *v*_*j*_ denote the feature vectors of protein node *i* and *j*, respectively. The least similar pairs were selected as the negative instances. As shown in **Supplementary Fig. 2**, there are five positive interactions (solid line) in the initial PPI network. For the positive pair *A* − *B*, we can select protein node *k* as a negative instance of *A* because it is the protein with the lowest similarity among all neighbors of node *B*.

#### Random Negative Sampling (RanNS)

In recommendation systems, Random negative sampling refers to uniformly sampling negative instances from the space of all answers ^51^. In our study, a random negative sampling strategy was designed for PPI prediction and compared with PopNS and SimNS. As an unbiased strategy, RanNS randomly selects a node that is not connected to it from the network and generates a negative instance.

### HGNNPIP model

#### The flowchart of the proposed HGNNPIP model

The flowchart of HGNNPIP is shown in **Figure 1**. The HGNNPIP framework consists of sequence embedding, network embedding, and MLP-based prediction model (**Figure 1A**). The *s*eq*uence embedding* module is developed to extract the global features of amino acid sequences (**Figure 1B**). The connectivity patterns involved in the PPI network are vectorized by our *network embedding* algorithm (**Figure 1C**).

**Figure 1.**
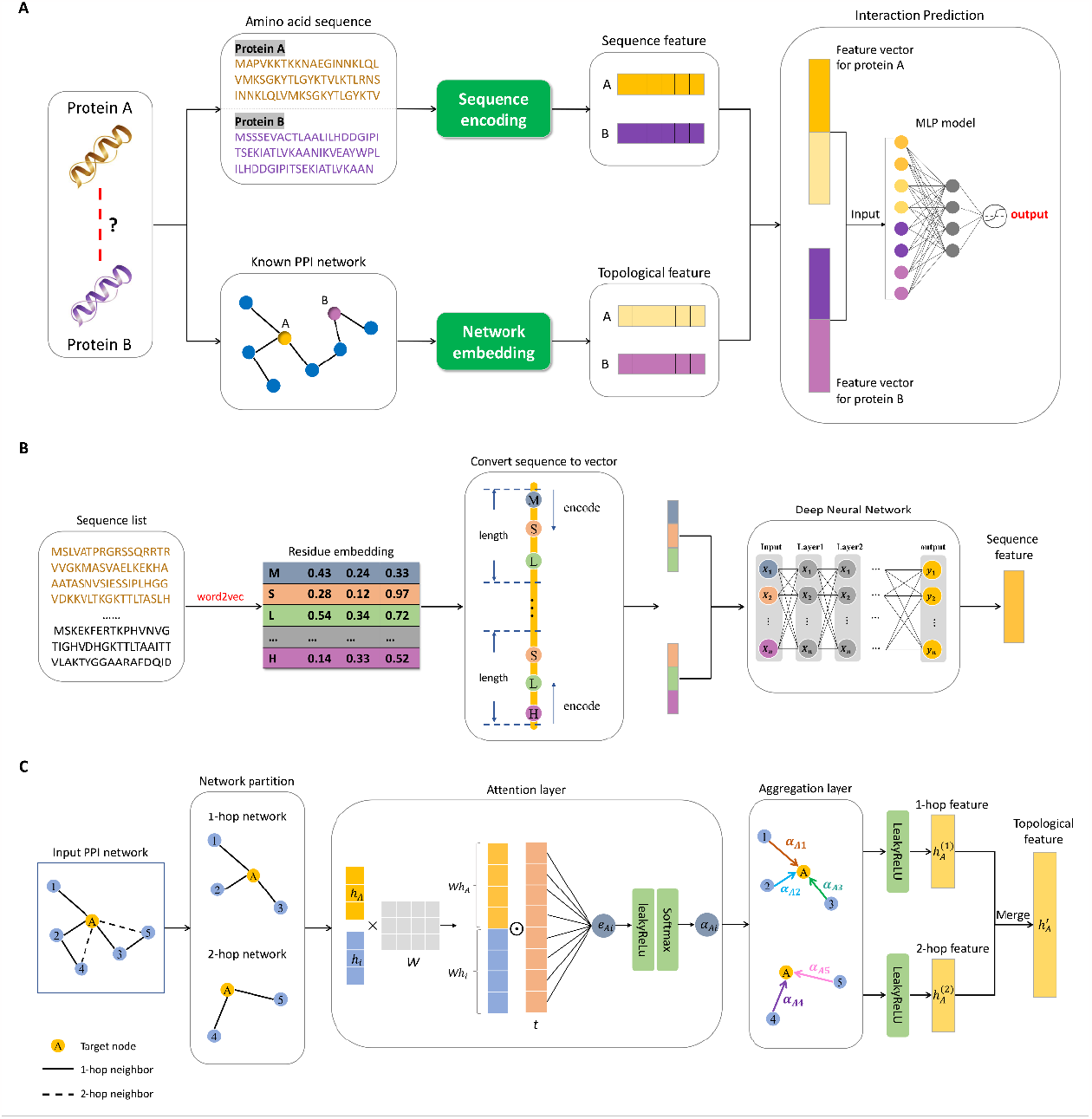
The procedure of HGNNPIP for PPI prediction. **(A)** The framework of HGNNPIP model; **(B)** Sequence embedding for residue feature representation; **(C)** Network embedding for connection property learning.

#### Sequence embedding strategy for residue feature representation

Sequence embedding process includes three stages: (i) amino acid vectorization using word2vec; (ii) protein sequence encoding; (iii) extracting abstract features from sequence encoding using Deep Neural Network (DNN) model.

##### Step 1: Amino acid vectorization using word2vec

As a successful word embedding technique in natural language processing ^52^, Word2vec can transform sentences into feature vectors with fixed dimensionality ^2,28^. An amino acid sequence of protein also can be considered as a ‘sentence’, which consists of 20 essential amino acids (‘words’). In this study, we used Word2vec model to transform each residue into a fixed dimensionality eigenvector representation (**Supplementary Fig. 3**). Through a sliding window, the center and context residues are segmented from the sequence and represented as the input and output of the neural network with one hot encoding ^2^. All the protein sequences in each dataset were adopted as a corpus to train the Word2vec model. The optimization of this model is to maximize the probability of the context word *S*[*i* ± 1] given a center word *S*[*i*]. We then obtained the embedding matrix *E* ∈ *R*^20×*r*^, in which the *j*-th row represents the feature vector of *j*-th amino-acid residue (*j* ∈ [1,20]). In another word, each amino-acid residue is embedded into a *r* −dimensional space.

##### Step 2: Protein sequence encoding

After generating the residue representation via Word2vec, we transformed each protein sequence into a vector with fixed dimensionality. Considering the fact that protein sequences may vary in length, we thus bidirectionally encoded the sequence from the head and tail ends. Given an amino acid sequence *S* = {*a*_1_, *a*_2_, …, *a*_*n*_}, two subsequences *S*_*start*_ = {*a*_1_, *a*_2_, …, *a*_*m*_} and *S*_*end*_ = {*a*_*n*−*m*+1_, *a*_*n*−*m*+2_, …, *a*_*n*_} are segmented from *S* . The parameter *n* and *m* denote the length of sequence *S* and a truncated subsequence (*S*_*start*_, *S*_*end*_), respectively. Therefore, the feature vector *X*_*start*_ and *X*_*end*_ for *S*_*start*_ and *S*_*end*_ can be described as follows (equation (2-3)).

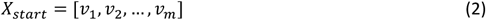

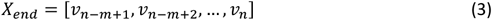

where *v*_*i*_ denotes the feature vector of the residue *a*_*i*_ generated by Word2vec model. We can easily find that the dimensionality of *X*_*start*_ and *X*_*end*_ are *r* ∗ *m* . Through the integration of subsequence representation (equation (4)), any sequence *S* is represented by a vector *X*^(0)^ and is sent into the DNN model.

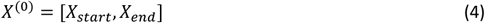

##### Step 3: Extracting deep features from sequence encoding using DNN

Considering that the dimensionality of *X*^(0)^ is high, we further designed a Deep Neural Network (DNN) model ^53^ to achieve dimension reduction (**Figure 1B**). Our DNN model includes multiple layers, and the output of *k*-th layer is defined as the following equation (5).

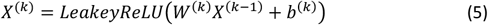

where *X*^(*k*−1)^ denotes the output of the layer *k* − 1. *W*^(*k*)^ and *b*^(*k*)^ represent the weight matrix of *k*-th layer and bias vector, respectively. LeakyReLu is a type of nonlinear activation function ^54^.

In this study, our DNN model includes 6 layers of neuro units. According to the length of the vector *X*^(0)^, the number of neuro units in the input layer is up to 2 ∗ *r* ∗ *m*. The size of the output layer is defined with parameter *z*. Furthermore, we constructed four hidden layers with 512, 256, 128, and 64 units. In addition, we incorporated BatchNorm and Dropout layers to enhance the ability of the model’s generalization. The parameter *r, m*, and *z* are automatically optimized by using Bayesian optimization ^55^.

### Network embedding strategy for connection property learning

As shown in **Figure 1C**, the network embedding module also includes three steps: (1) network partition; (2) attention calculation; (3) topological feature extraction.

#### Step 1: Network partition

Here, we take protein *A* as an example to illustrate the procedure of network embedding. First, we need to calculate the directed path from the vertex *V*_*A*_ to any other nodes. Then, the 1-hop and 2-hop networks associated with the node *A* are extracted from the initial PPI network. In order to drive a graph neuron network, each node needs to be initialized first ^56^. In this study, we combine embedded sequence vectors with connectivity information to assign the initial state of the nodes. The node *A* can be defined as follows:

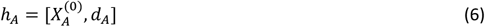

where 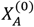 denotes the sequence embedding vector obtained from Eq. (4). *d*_*A*_ represents the degree of node *A*. Similarly, any vertex *V*_*i*_ (protein *i*) in the 1-hop or 2-hop neighbors of a vertex *V*_*A*_ can be initialized as:

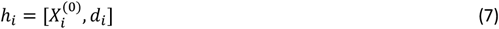

where 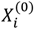and *d*_*i*_ denote the sequence vector and degree of vertex *V*_*i*_.

#### Step 2: Attention calculation

To assign a weighting factor (*α*_*Ai*_) to each connection, our model calculates the similarity between a vertex *V*_*A*_ and any (*V*_*i*_) of its 1-hop or 2-hop neighbors. This weighting factor is also called ‘attention’ and reflects the importance of the protein *i*. The similarity (*e*_*Ai*_) between vertex *V*_*A*_ and *V*_*i*_ is defined by the following equation (8):

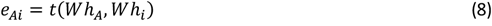

where *W* and *t* are coefficient matrix (*W* ∈ *R*^*F*′×*F*^, *t* ∈ *R*^2*F*′×1^). Here, the lengths of *h*_*A*_ and *h*_*i*_ are equal to *F* . While the lengths of *Wh*_*A*_ and *Wh*_*i*_ are both *F*^′^ . Finally, the weighting factor *α*_*Ai*_ is determined by equation (9):

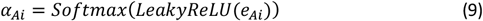

#### Step 3: Topological feature extraction

After obtaining all the weighting factors for 1-hop and 2-hop networks, the topological feature of protein *A* can be represented through a neighborhood aggregation scheme. Firstly, the 1-hop feature of protein *A* is defined by equation (10).

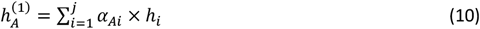

Similarly, the 2-hop feature of protein A is defined by equation (11).

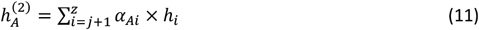

where the number of 1-hop and 2-hop neighbors are *j* and *z* − *j*. Finally, the topological feature of protein *A* is achieved by merging the vector 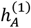and 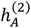:

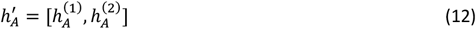

### PPI prediction with Binary Classifier

Based on the above description, protein *A* can be represented as a concatenation vector *p*_*A*_ (equation (13)):

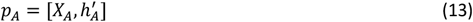

where *X*_*A*_ and 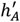 denote the embedding vector of sequence and topological feature, respectively. Given another protein *B* with feature vector 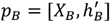, the MLP classifier determines whether it has a potential interaction with *A*. As shown in **Figure 1A**, the size of the input layer is equal to the sum of the lengths of *p*_*A*_ and *p*_*B*_. In addition, we use one output node to return the probability that any pair of proteins may interact. In this study, we used 0.5 as the probability cutoff to assign labels ^30,34,35^.

#### Cross-validation

To perform model selection, 5-fold cross-validation was set up in this study. Under this setting, each dataset is equally divided into five folds and each part has an equal chance to train and test the models. Therefore, the same process is independently repeated 5 times, and 5 optimized models are obtained. Finally, averaging over 5 splits yielded an estimate of model performance.

#### Performance metrics of prediction model

We combined eight evaluation metrics to assess the prediction performance of the models, including Accuracy, Precision, Recall, Specificity, F1 score, Matthews Correlation Coefficient (MCC), Area Under the Receiver Operating Characteristic (AUROC), and Average Precision (AP) ^2^. Higher values in all these metrics indicate better performance. The definition of the first six metrics is presented as follows (equation (14-19)):

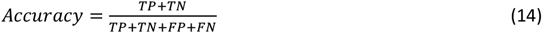

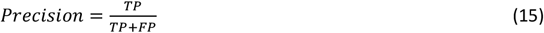

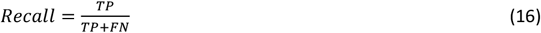

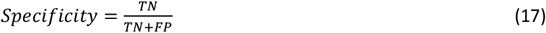

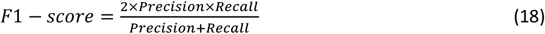

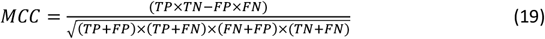

where four indicators *TP, TN, FP, FN* denote True Positive, True Negative, False Positive, and False Negative, respectively. Moreover, two area-associated metrics are always used to evaluate the model performance ^26^. AUROC illustrates how much the model is capable of distinguishing between classes ^57^. In addition, AP is also a popular metric to summarize the Precision-Recall Curve to one scalar value ^58^.

### Evaluation metric of negative sampling strategies

To analyze the impact of various negative sampling strategies on model outcome, we defined a new metric (*α*) to reflect the false positive rate (FFR) and Recall (equation (20)).

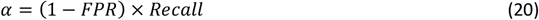

As shown in **Eq. (20)**, a better strategy should first guarantee the generated negative instances do not cause high false positives while maintaining the high true positive rate.

#### The effect of the ratio of negative to positive samples on model outcome

In previous studies, many researchers tend to construct balanced datasets, that is, the number of positive and negative instances is the same or close ^2,35,37^. However, there are significantly more potential noninteracting pairs than interacting pairs in any dataset. In this study, we conducted an in-depth analysis of the impact of the ratio (*k*) of negative to positive samples on the model outcome.

#### Comparisons with existing algorithms

In the comparative study, our HGNNPIP model was compared with four SOTA models, including DeepTrio ^27^, PIPR ^29^, DeepFE-PPI ^28^, and GAT ^59^. All the comparisons were implemented using 5-fold cross-validation on six PPI datasets. The first three methods all achieve PPI prediction by combining protein sequence information with deep learning models. In particular, both DeepTrio and PIPR are established on CNN model, while DeepFE-PPI depends on a DNN architecture. GAT, is a graph neural network model originally proposed by Velickovic et al., for protein-protein interaction.

#### Simulation environment

The code of the HGNNPIP model was developed and debugged using Tensorflow 2.7.0 and Python 3.7.0 under the environment with Ubuntu 18.04, GPU 3090 (24G), and 512G RAM.

#### Plant Preparation

Seeds of wild-type (*Oryza sativa* L. ZH11) and transgenic plants (*Oseds1*) were surface sterilized with 10% NaClO, washed with ddH_2_O, and imbibed in 4°C overnight. The imbibed seeds were germinated on soil at 28°C. The germinal seedlings at 3 days were transformed into pots filled with nutrient soil, and then grown in a growth chamber under the conditions of 14-h day, 28°C, 80% relative humidity (RH) followed by 10-h night, 26°C, 80% RH.

#### Bacterial inoculation

For the Xanthomonas oryzae pv. oryzae (*Xoo)* infection assay, bacterial strain PXO99A was grown on nutrient broth agar (NA) medium at 28°C for 2 days, bacterial pellets were washed twice and resuspended in sterile water to generate inoculum suspensions with an optical density of OD_600_ = 0.5. The *Xoo* suspension was directly inoculated on leaves of two-week-old rice seedlings by infiltration with needleless syringes with three sites per leaf. Three leaves were collected as one sample for RNA extraction.

#### RNA sequencing and data preprocessing

*Xoo* infected leaves at 48 hour-post-infiltration were collected with three leaves at each time point as one biological replicate. Three biological replicates of samples were collected. RNA-seq analysis was performed by Majorbio BioTech Co. (Shanghai, China). The Illumina Novaseq 6000 platform was used to construct RNA libraries and generate reads of 300 bp long paired-end (Illumina, San Diego, CA, USA). The read number of each gene was transformed into FPKM (fragments per kilobase of exon model per million mapped reads). After quality control of the raw data, the clean data were compared with the Oryza_sativa reference genome (IRGSP-1.0) using TopHat ^60^ (v2.1.1) to obtain mapped data, which were used for transcript assembly and expression calculation. The expression levels of genes and transcripts were quantified using RSEM ^61^ (v1.3.3). After obtaining read counts of genes, differentially expressed genes were identified using the DESeq2 ^62^ (V1.24.0) and the identification criteria were set as adjusted P-value <0.05 and |log_2_FC|≥ 0.8.

#### qPCR validation

Total RNA was extracted with TRIzol reagent (Thermo Fisher Scientific, 15596026) from leaf tissues. Extracted RNA was treated with gDNA wiper, and 1 μg of total RNA was reverse-transcribed using HiScript II Reverse Transcriptase (Vazyme, R223) with Random primers/Oligo(dT)_23_ primer mix. Real-time DNA amplification was conducted using Bio-Rad iQ5 optical system software (Bio-Rad). OsActin was used as the internal control. The primers used for qRT-PCR are listed in **Supplementary Table 1**.

## 3. Results

### Bias in data distribution may cause a high false positive rate

In previous works, computational scientists tried to access various public databases to obtain the interacting data (positive instances), which were mostly validated by experiments. To construct balanced datasets, they manually generated the same number of negative instances through subcellular localization ^2,35,37^. However, these negative PPIs are only associated with a small portion of proteins (from 7.58% to 47.22%), indicating serious data distribution bias on most benchmarking datasets (**Table 1**).

To determine the impact of data distribution bias on false positives in model prediction, we created 2 testing sets for each PPI dataset, named as ‘NTS1’ and ‘NTS2’, each containing 100 randomly selected negative instances based on subcellular localization information. All the instances in ‘NTS1’ are associated only with the proteins in the original dataset that established negative PPIs via subcellular localization (5^th^ column in **Table 1**). However, the proteins used to construct ‘NTS2’ have only participated in positive PPIs. HGNNPIP and other SOTA models were all trained on each original dataset and then tested on ‘NTS1’ and ‘NTS2’ datasets. **Table 2** shows that all the computational models had high false positives for their predictions on ‘NTS2’ dataset (at least 90% of negative samples are predicted as positive). On the contrary, the false positive rate on NTS1 is significantly lower. The above analysis indicates that the false positives are mainly due to the bias in data distribution rather than the model design.

### The effect of negative sampling strategy on model outcome

Since subcellular localization can lead to high false positive rates and low generalization, we introduced three potential negative sampling strategies (PopNS, SimNS, RanNS) and evaluated their effects on model outcomes. This analysis was implemented on the dataset of *S. cerevisiae*. As shown in **Table 3**, all three negative sampling strategies significantly reduce the false positive rate (FPR) rather than subcellular localization, while they also lead to lower recall as the diversity of negative samples.

In this study, we defined a metric to minimize false positive and false negative rates, simultaneously. From **Table 3**, we can easily find that random negative sampling (RanNS) is the best solution to control false positives and negatives. In further analysis, we chose RanNS to correct all six benchmarking datasets.

### Determine the optimal ratio of negative to positive instances

In this subsection, we explore the optimal ratio of negative to positive instances when constructing a PPI dataset for training a predictive model. HGNNPIP and 4 SOTA models were tested on the dataset *S. cerevisiae* with the value of *k* from 1 to 5. As shown in **Figure 2**, both FPR and Recall of each model obviously decrease as *k* increases. Moreover, HGNNPIP returns the best *α* when *k* is equal to 2. Similarly, PIPR received the optimal *α* if *k* = 3. For the other three models, *k* equal to 1 seems to be appropriate.

**Figure 2.**
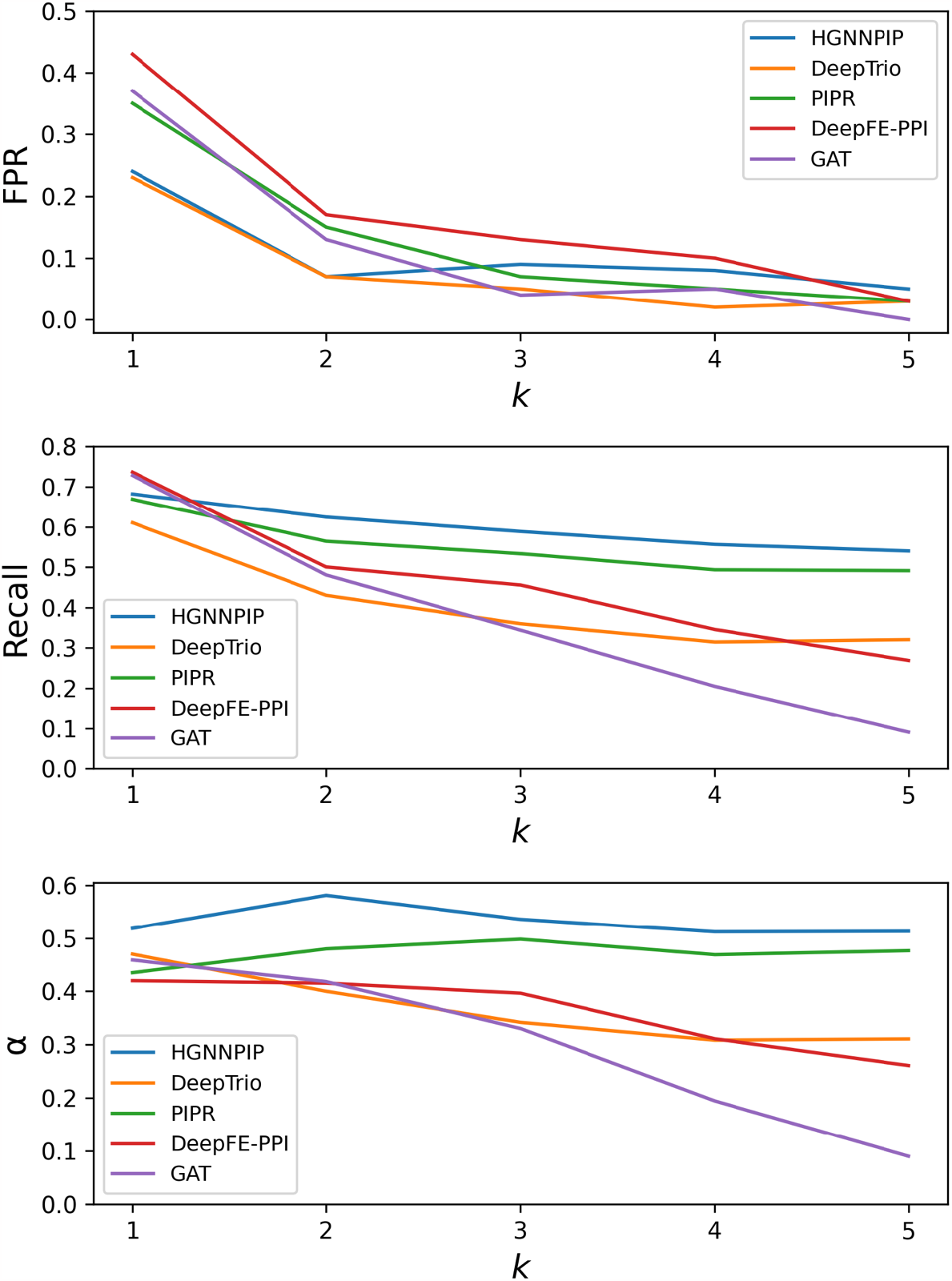
The effect of parameter *k* on model outcome. *k* is defined as the ratio of negative to positive samples.

In addition, we also examined 8 performance metrics to determine how the parameter *k* affects model outcome. In **Table 4**, we can clearly see that the HGNNPIP model with the optimal *k* (*k* = 2) not only controls false positives and negatives very well, but also guarantees the prediction accuracy.

### Dataset reconstruction using Random Negative Sampling

Based on the above description, we generated 2 folds of negative instances (*k* = 2) using RanNS based on the positive PPIs shown in **Table 1** and reconstructed all six datasets. Since the training set for each species was adjusted, we tested the false positive rate on testing set NTS2. As shown in **Table 5**, the false positive rates of all prediction models have been significantly reduced after reconstructing the PPI datasets through negative sampling strategy. Therefore, the proposed Random Negative Sampling (RanNS) strategy is suitable for general PPI prediction models.

### Parameter selection for HGNNPIP modeling

After correcting the distribution of negative samples and reconstructing datasets, we then completed parameter selection to optimize HGNNPIP modeling. Overall, there are 7 key parameters need to be determined, which are associated with feature representation and neural network construction. The details of the above parameters can be seen in **Table 6**.

Here, we mainly considered ‘Accuracy’ as the representative evaluation metric and used the Bayesian method ^63^ to implement parameter optimization on the *S. cerevisiae* dataset. As shown in **Table 6**, the whole searching process includes 15 rounds, and the best parameter set was obtained in the 6^th^ round. For simplicity, the above optimal parameters were transferred when HGNNPIP was applied to other datasets.

### Improving model performance by combing sequence and network information

To analyze the contributions of feature fusion in HGNNPIP, we designed an ablation study, in which we tested two baseline models (with sequence feature, or network feature) and the integrated HGNNPIP model on six corrected PPI datasets. From **Figure 3**, we can see that the integrative model significantly outperforms the model with only sequence or network features on 7 out of 8 metrics. In addition, the predictive model with network features provides better performance than sequence features in six metrics, including Accuracy, Recall, F1-score, MCC, AUROC, and AP (**Table 7**). Therefore, integrating features of amino acid sequences and network connections potentially improves model performance.

**Figure 3.**
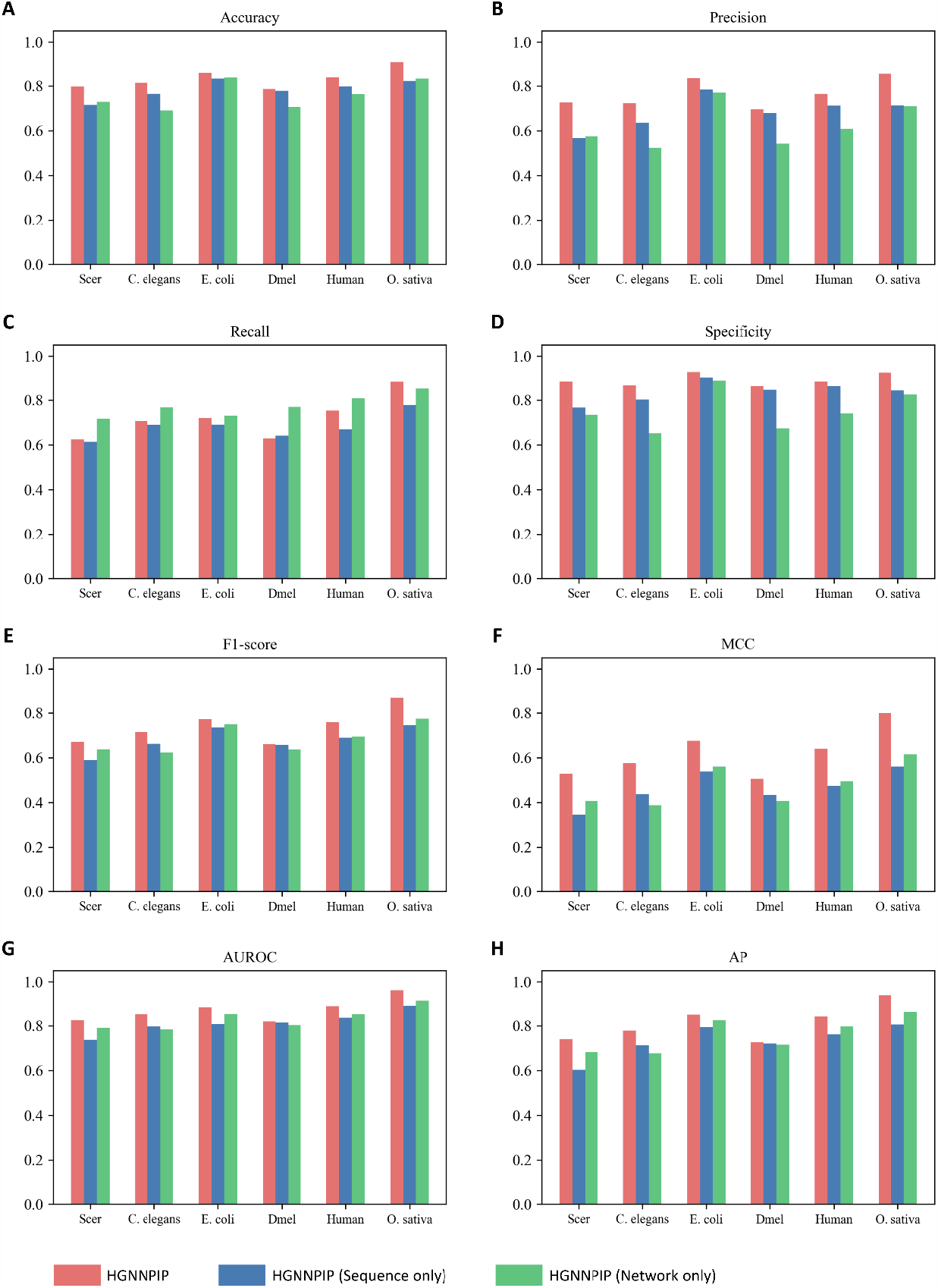
The ablation experiments of HGNNPIP model on six PPI datasets.

### HGNNPIP outperforms the SOTA algorithms

To verify the outstanding performance, we compared the proposed HGNNPIP model with the other four SOTA methods (DeepTrio, PIPR, DeepFe-PPI, and GAT) on all six reconstructed datasets (**Figure 4**). Overall, HGNNPIP significantly outperforms all other SOTA models on four datasets, including *S. cerevisiae, C. elegans, Human, O. sativa*. Moreover, HGNNPIP provides the best accuracy, recall, MCC, and AUROC on all six species, indicating its great prediction power for unbalanced datasets. Among four SOTA models, DeepTrio and PIPR perform better. Taken together, HGNNPIP is superior to the existing methods in terms of accuracy and generalization (**Table 8**).

**Figure 4.**
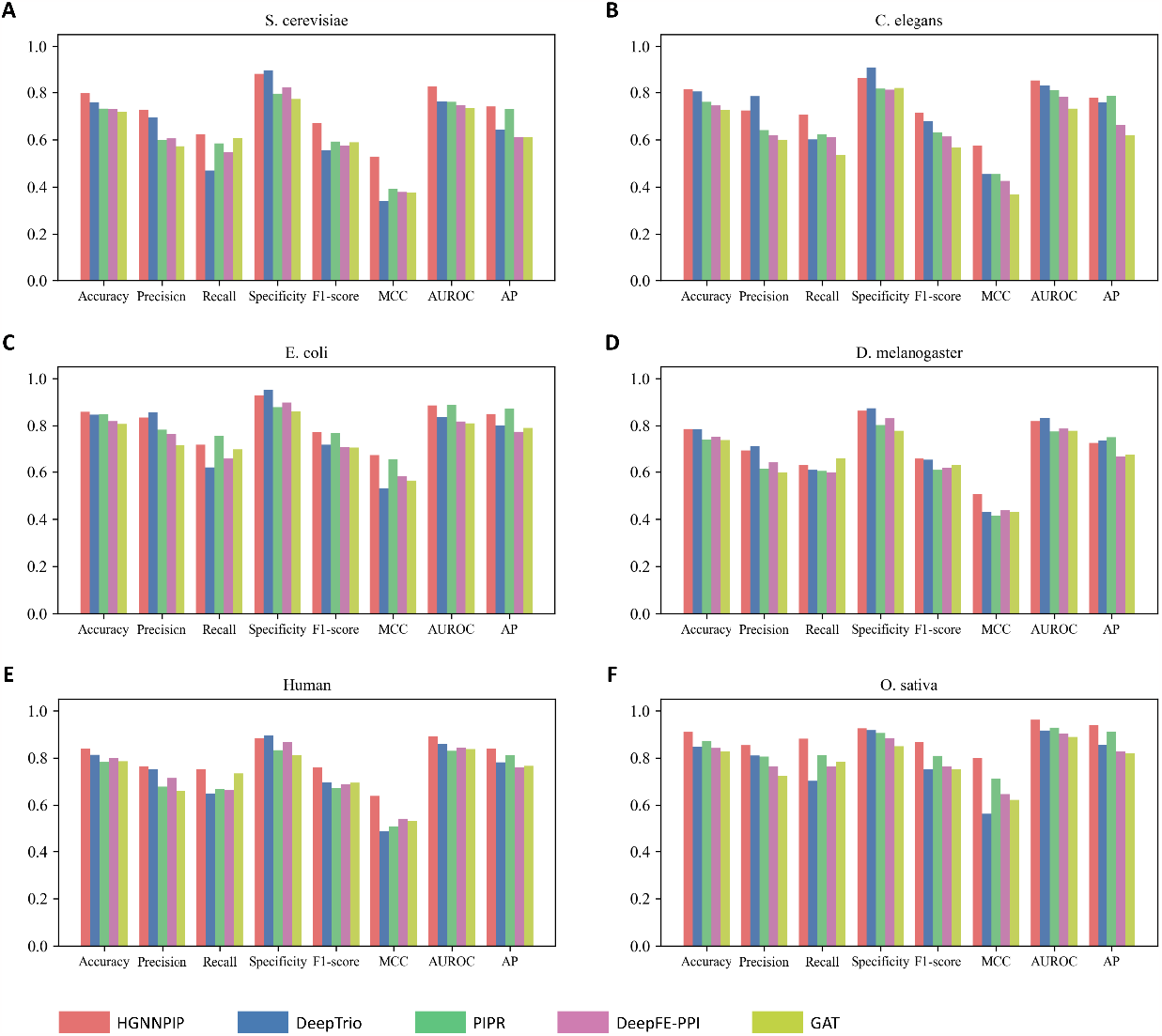
Comparisons of HGNNPIP with four SOTA models on the PPI datasets across six species, including **(A)** *S. cerevisiae*; **(B)** *C. elegans*; (**C)** *E. coli*; **(D)** *D. melanogaster*; **(E)** *Human*; **(F)** *O. sativa*

### Applying HGNNPIP model to the study of Oryza sativa

In this section, we conducted a case study of *Oryza sativa* to determine how HGNNPIP model identifies potential molecular contacts in diseased rice.

#### 1) Identifying the interacting partners of OsEDS1 in infected rice

Previous studies indicated that OsEDS1 signaling may be involved in positively regulating disease resistance in rice; however, the underlying mechanisms remain unclear ^64^. Here, we integrated the proposed HGNNPIP model with our transcriptome data (3 mutant vs. 3 control) of rice to predict the possible interacting partners of OsEDS1. Our DEG analysis ^3^ screened out 1204 up-regulated and 903 down-regulated genes (**Supplementary Fig. 4**). By overlapping the *O. sativa* PPI network (**Table 1**), we obtained 216 differential expressed proteins for further prediction. **Figure 5A** shows 30 interacting partners of OsEDS1 with high confidence predicted by the HGNNPIP model. 15 representative proteins with green color were reported by literature ^65,66^. The remaining proteins (marked in red) were identified for the first by us (**Table 9**). Furthermore, we implemented *In vivo* experiments to validate the potential interactions predicted by the HGNNPIP model. PCR analysis revealed that the transcriptional level of six proteins was significantly altered in the *Xoo*-infected rice with *Oseds1* mutant (**Figure 5B**). Except for OsHOX1, the other five proteins (e.g., OsWRKY72, OsWRKY45, *etc*.) were all down-regulated in the rice with *Oseds1 mutant*. The above findings suggest that these six proteins are involved in the OsEDS1-mediated regulation of disease resistance and the crosstalk with abiotic stress and development in rice.

**Figure 5.**
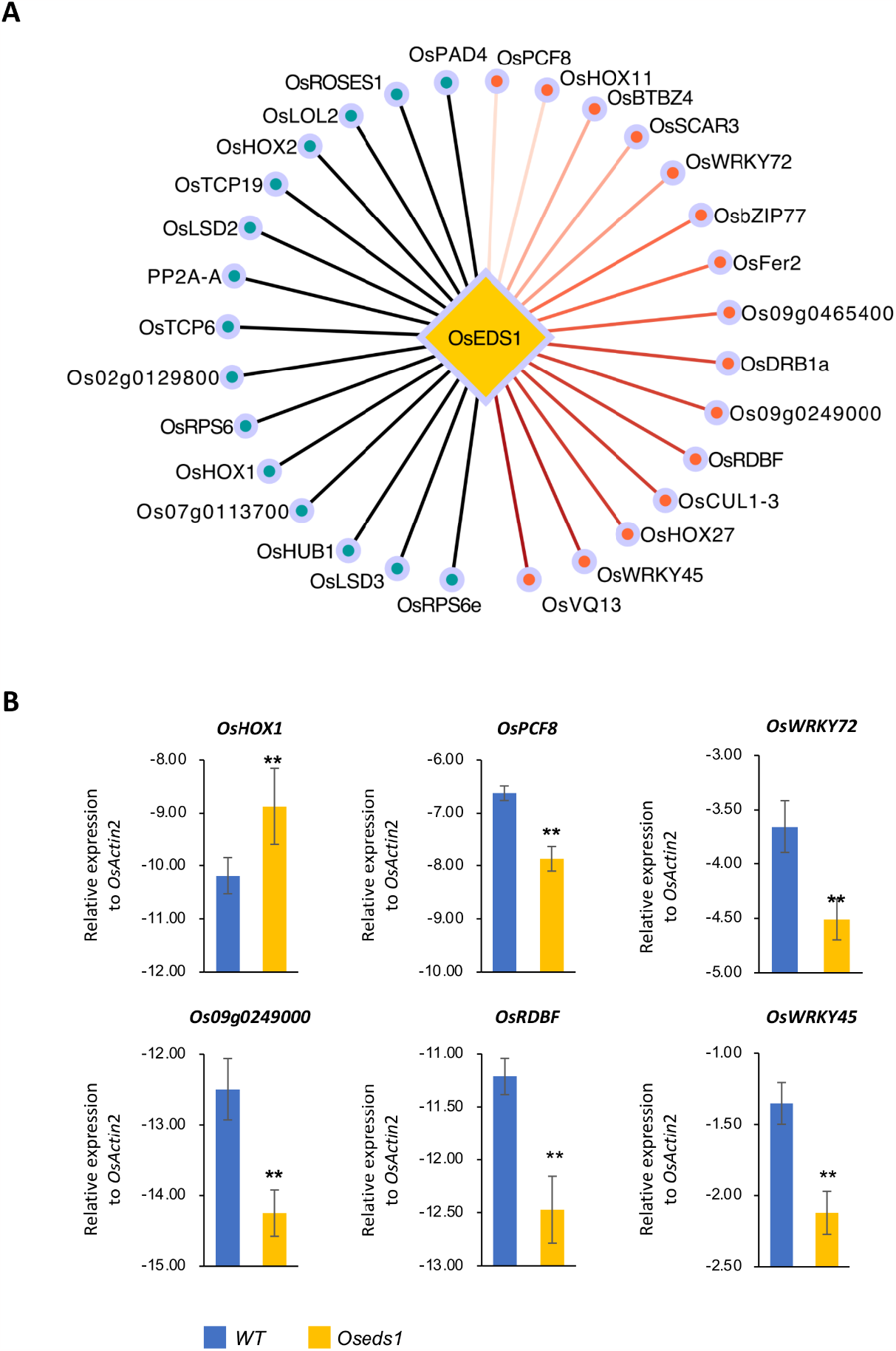
Prediction and validation of OsEDS1-associated interacting proteins. **(A)** The OsEDS1-specific network predicted by HGNNPIP; **(B)** qPCR validations on six potential factors.

#### 2) Predicting the potential targets of SCRE4 in rice false smut

Previous studies revealed that the *Ustilaginoidea virens-*secreted effector *SCRE4* inhibits the expression of the positive immune regulator OsARF17, thereby suppressing rice immunity ^67^. However, the biological function of *SCRE4* has not been fully understood. In this study, we applied the HGNNPIP model on our transcriptome data from the rice infected by *Ustilaginoidea virens* (3 infected vs. 3 control) to explore potential targets of *SCRE4*. Similarly, pathway analysis was implemented using DAVID ^68^. 24 proteins were identified to associate with pathogen-host interactions (**Supplementary Fig. 5**). HGNNPIP provides confidence scores for *SCRE4* interactions with these 24 host proteins (**Table 10**). As shown in **Table 10**, OsRLCK33 and OsCML11 appear to be two factors with high possibilities. Based on model predictions, we finally inferred the *SCRE4*-specific network, which consists of three functional modules (**Figure 6**). Except for the protein OsARF17 mentioned above, both OsRLCK33 and OsCML11 also seem to serve as important hub nodes in regulating rice development and defense response.

**Figure 6.**
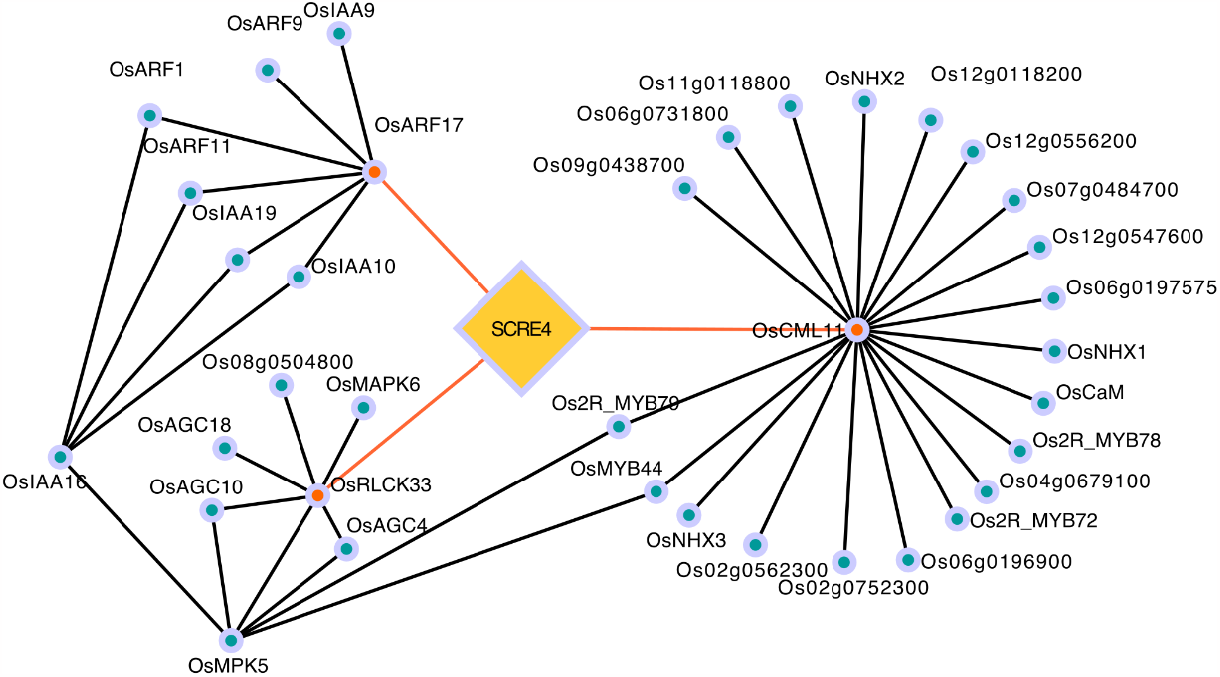
The SCRE4-specific network predicted by HGNNPIP.

## 4. Discussion

A deep understanding of PPIs can provide global insights into cellular organization, genome function, and genotype-phenotype relationships in various species ^69^. Increasingly, computational algorithms are being developed to discover previously unrecognized PPIs and, thereby, improve the comprehensiveness of the interactome map ^1^. In this study, we proposed a Hybrid Graph Neural Network model (HGNNPIP) for Protein-protein interaction prediction. By combining the features extracted from amino acid sequences and the connection properties of known PPIs, HGNNPIP achieved excellent performance across different species. The comparative analysis further demonstrated that the HGNNPIP model is superior to the other four SOTA models in PPI prediction.

### Excellent performance of HGNNPIP is mainly attributed to feature fusion

We fully extracted the intrinsic properties of proteins to determine the probability of interacting relationships. Combining sequence and network information can significantly improve the prediction performance. According to **Table 8**, the integrated model can increase the Accuracy, AUROC, and AP by approximately 4.54-8.63%, 3.50-6.21%, and 4.54-8.63%. Unlike many deep learning models reported in the previous studies ^27,28,30-32,70^, HGNNPIP implements the prediction with a simple classifier MLP. It indicates that the computational cost of our algorithm will not be high. Compared with the other three general classifiers (SVM, Random Forest, Linear regression) on dataset *S. cerevisiae*, we found that MLP is the best choice for inclusion in the HGNNPIP framework (**Table 11**).

### Subcellular localization induces a high false positive rate of machine learning models

As mentioned in the section of “**Materials and Methods**”, most computational scientists tend to generate negative pairs by randomly pairing proteins with different subcellular localization ^2,34,35,37^. Therefore, these representative benchmark PPI datasets are manually curated with an equal number of positive and negative samples. The information regarding the subcellular localization of proteins can be searched from the Swiss-Prot database ^40^. However, the negative instances generated in this way may cause bias in the data distribution, i.e., only a small portion of proteins are associated with these non-interacting examples (**Table 1**). Training a predictive model with a biased dataset may obtain good fitting accuracy (overfitting) but low generalization ^71^.

### Dataset construction based on positive instances for model training

In this study, we proposed three possible strategies for negative instance generation, including Popularity-biased Negative Sampling (PopNS) ^42-44^, Similarity-biased Negative Sampling (SimNS) ^48-50^, and Random Negative Sampling (RanNS) ^51^. The underlying assumption of these methods is that a protein pair can be determined as a negative instance if there is currently no evidence to prove that they may interact. Our analysis demonstrated that data correction with the above strategies can sharply reduce the false positive rate for a general predictive model. RanNS is the best choice for generating high-quality negative instances before training predictive models. Actually, RanNS also provides better model performance than PoPNS and SimNS in the recommendation systems ^45,50^. Taken together, the PPI datasets constructed by RanNS ensure excellent performance of predictive models with high fitting accuracy and strong generalization ability.

Furthermore, the case study on *Oryza sativa* demonstrated that our HGNNPIP has great potential to reveal the molecular mechanisms. OsWRKY45 was reported up-regulated during *Xoo* infection in rice, whereas overexpression of OsWRKY45 leads to enhanced resistance against bacterial pathogens in *Arabidopsis* ^72^. In our study, we noticed that OsWRKY45 was down-reduced in the Oseds1 mutant condition, indicating that OsWRKY45 may work as a downstream factor of OsEDS1 and is required for OsEDS1-mediated disease resistance. OsWRKY72 was reported as a negative regulator of *Xoo* resistance by suppressing jasmonic acid biosynthesis ^73^. These findings illustrated the possible role of WRKY transcription factors in OsEDS1-mediated immunity. EDS1 has been reported to regulate SA- and ABA-mediated stomatal immunity against bacterial pathogens through WRKY transcription factor in *Nicotiana benthamiana* and Arabidopsis ^74^. OsPCF8, belonging to the TCP transcription factor family, is required for cold tolerance in rice ^75^. Interestingly, the *eds1* mutant showed better performance under cold stress in *Arabidopsis* ^76^. Furthermore, OsRDBF and OsHOX1 are closely related with gain filling and tiller angle in rice, respectively ^77,78^. These results suggesting that OsEDS1 may be also important for the crosstalk between biotic-abiotic stresses and defense-growth. However, the detailed mechanism still needs to be investigated in the near future.

Limitations still exist in the current study. Integrating protein structure information may improve the prediction performance. Parameter optimization on dataset *S. cerevisiae* is not necessarily optimal for other datasets. Except for the SOTA models mentioned in this work, it is unknown whether HGNNPIP is better than other existing methods. The dynamics of protein-protein interactions are also valuable to be studied. In future work, transgenic plant overexpressing or knock-out target genes will be generated to further validate the function of putative function in SCRE4-mediated interaction between host and pathogen.

## Data availability

The public datasets we used were from refs. ^35,37^, and the usages are fully illustrated in **Methods**. All six processed PPI datasets are placed in the link: doi.org/10.6084/m9.figshare.24763902. The researchers can also download the mirror data from the link: http://cdsic.njau.edu.cn/data/PPIDataBankV2.0. All the transcriptome profiles of *Oryza sativa* can be accessed through: GSE248248.

## Code availability

An open-source implementation of the HGNNPIP model is available at https://github.com/chilutong/HGNNPIP.

## Author’s contributions

Lutong Chi: Data analysis and modeling, Coding, Writing.

Jinbiao Ma: Sample collection, Experimental validation, Writing.

Yingqiao Wan: Experimental validation.

Yang Deng: Software.

Yufeng Wu: Resources.

Xiaochen Cen: Data preprocessing.

Xiaobo Zhou: Review & editing.

Xin Zhao: Review & editing.

Yiming Wang: Conceptualization, Funding acquisition, Review & editing.

Zhiwei Ji: Supervision, Project administration, Funding acquisition, Conceptualization, Methodology, Review & editing.

## Competing interest statement

The authors declare that they have no known competing financial interests or personal relationships that could have appeared to influence the work reported in this paper.

## Acknowledgements

The authors thank Dr. Weiling Zhao at UNC Chapel Hill for her valuable writing advice. This work was supported by the Fundamental Research Funds for the Central Universities (No. KYCXJC2023001, No. KJJQ2023001), Natural Science Foundation of Jiangsu Province (No. BK20211210, BK20220085), and Agricultural Science and Technology Innovation Foundation of Jiangsu Province (No. CX (23) 3125). This work was partially supported by the startup award of new professors at Nanjing Agricultural University (No. 106/804001).

## Notes

http://cdsic.njau.edu.cn/data/PPIDataBankV2.0

### Competing Interest Statement

The authors have declared no competing interest.

