## Supplementary material for "HGNNPIP: A Hybrid Graph Neural Network framework for Protein-protein Interaction Prediction": Tables

**Table 1.** Summary of six PPI datasets used in this study. Negative PPIs are originally generated by subcellular localization. The first five datasets are directly downloaded from literatures.

| Datasets | Proteins | Positive PPIs | Negative PPIs | Proteins in Negative PPIs | Rate | PMID |
|---|---|---|---|---|---|---|
| *S. cerevisiae* | 2526 | 5594 | 5594 | 1193 | 47.22% | 20500905 |
| *C. elegans* | 2638 | 4030 | 4030 | 200 | 7.58% | 20500905 |
| *E. coli* | 1832 | 6954 | 6954 | 564 | 30.75% | 20500905 |
| *D. melanogaster* | 7059 | 21975 | 21975 | 635 | 8.99% | 20500905 |
| *Human* | 10364 | 36630 | 36480 | 2184 | 23.10% | 20698572 |
| *O. sativa* | 7920 | 33876 | 33876 | 1183 | 14.94% | string-db.org |

**Table 2**. The predictions of five models on two independent testing sets. Both NTS1 and NTS2 contain 100 negative instances. The ratio of positive to negative is equal to 1:1 ($k = 1$).

| Dataset | Method | False positive rate for NST1 | False positive rate for NST2 |
|---|---|---|---|
| *S. cerevisiae* | HGNNPIP | 0.14 | 1 |
| | DeepTrio | 0.21 | 1 |
| | PIPR | 0.24 | 0.84 |
| | DeepFE-PPI | 0.15 | 0.96 |
| | GAT | 0.13 | 0.85 |
| *C. elegans* | HGNNPIP | 0.06 | 1 |
| | DeepTrio | 0.04 | 1 |
| | PIPR | 0.04 | 0.98 |
| | DeepFE-PPI | 0.07 | 1 |
| | GAT | 0.09 | 0.89 |
| *E. coli* | HGNNPIP | 0.06 | 0.96 |
| | DeepTrio | 0.10 | 0.98 |
| | PIPR | 0.12 | 0.97 |
| | DeepFE-PPI | 0.07 | 0.82 |
| | GAT | 0.11 | 0.90 |
| *D. melanogaster* | HGNNPIP | 0.06 | 1 |
| | DeepTrio | 0.05 | 1 |
| | PIPR | 0.35 | 1 |
| | DeepFE-PPI | 0.05 | 1 |
| | GAT | 0.03 | 0.95 |
| *Human* | HGNNPIP | 0.10 | 0.98 |
| | DeepTrio | 0.07 | 0.98 |
| | PIPR | 0.33 | 0.95 |
| | DeepFE-PPI | 0.16 | 0.97 |
| | GAT | 0.10 | 0.92 |
| *O. sativa* | HGNNPIP | 0.07 | 1 |
| | DeepTrio | 0.05 | 1 |
| | PIPR | 0.21 | 0.91 |
| | DeepFE-PPI | 0.10 | 0.98 |
| | GAT | 0.09 | 0.96 |

**Table 3.** The effect of negative sampling strategy on model outcome. The ratio of positive to negative is equal to 1:1 ($k = 1$). Analysis is implemented on the original dataset *S. cerevisiae.* FPR denotes false positive rate.

| **Metrics** | **Negative Sampling method** | **HGNNPIP** | **DeepTrio** | **PIPR** | **DeepFE-PPI** | **GAT** |
|---|---|---|---|---|---|---|
| FPR | Subcellular localization | 1 | 1 | 0.84 | 0.96 | 0.85 |
| | Popularity-biased Negative Sampling | 0.4 | 0 | 0.45 | 0.41 | 0.55 |
| | Similarity-biased Negative Sampling | 0.37 | 0 | 0.45 | 0.56 | 0.47 |
| | Random Negative Sampling | 0.24 | 0.23 | 0.35 | 0.43 | 0.37 |
| Recall | Subcellular localization | 0.9458 | 0.8928 | 0.8850 | 0.9219 | 0.8922 |
| | Popularity-biased Negative Sampling | 0.5426 | 0 | 0.5800 | 0.4361 | 0.4478 |
| | Similarity-biased Negative Sampling | 0.6774 | 0.0678 | 0.7017 | 0.6917 | 0.5968 |
| | Random Negative Sampling | 0.6821 | 0.6103 | 0.6690 | 0.7366 | 0.7283 |
| $\alpha$ | Subcellular localization | 0 | 0 | 0.1416 | 0.036 | 0.1338 |
| | Popularity-biased Negative Sampling | 0.3256 | 0 | 0.319 | 0.2573 | 0.2463 |
| | Similarity-biased Negative Sampling | 0.4267 | 0.0678 | 0.315 | 0.3043 | 0.3163 |
| | Random Negative Sampling | **0.5183** | **0.4455** | **0.4349** | **0.3167** | **0.4588** |

**Table 4.** The effect of the ratio of negative to positive samples on model outcome. All analysis is implemented on the dataset *S. cerevisiae*.

| $k$ | Model | Accuracy | Precision | F1-score | MCC | AUROC | AP | Recall | FPR | $\alpha$ |
|---|---|---|---|---|---|---|---|---|---|---|
| 1 | HGNNPIP | 0.7252 | 0.748 | 0.7136 | 0.307 | 0.77 | 0.7619 | 0.6821 | 0.24 | **0.5184** |
| | DeepTrio | 0.6979 | 0.7337 | 0.6663 | 0.2498 | 0.744 | 0.747 | 0.6103 | 0.23 | 0.4699 |
| | PIPR | 0.6658 | 0.6655 | 0.6673 | 0.3315 | 0.7236 | 0.7041 | 0.6690 | 0.35 | 0.4349 |
| | DeepFE-PPI | 0.6917 | 0.6827 | 0.7086 | 0.2754 | 0.7509 | 0.7258 | 0.7366 | 0.43 | 0.4199 |
| | GAT | 0.6676 | 0.6494 | 0.6866 | 0.3376 | 0.7187 | 0.7104 | 0.7283 | 0.37 | 0.4588 |
| 2 | HGNNPIP | 0.7964 | 0.7268 | 0.6716 | 0.5288 | 0.826 | 0.6716 | 0.6246 | 0.07 | **0.5808** |
| | DeepTrio | 0.7504 | 0.7162 | 0.5375 | 0.3155 | 0.7662 | 0.6531 | 0.4302 | 0.07 | 0.4001 |
| | PIPR | 0.7355 | 0.6190 | 0.5904 | 0.3966 | 0.7640 | 0.7378 | 0.5644 | 0.15 | 0.4797 |
| | DeepFE-PPI | 0.7367 | 0.6328 | 0.5589 | 0.3801 | 0.7508 | 0.6091 | 0.5004 | 0.17 | 0.4153 |
| | GAT | 0.7385 | 0.6105 | 0.5379 | 0.319 | 0.7636 | 0.6073 | 0.4807 | 0.13 | 0.4182 |
| 3 | HGNNPIP | 0.8288 | 0.6492 | 0.6174 | 0.4732 | 0.799 | 0.658 | 0.5886 | 0.09 | **0.5356** |
| | DeepTrio | 0.7994 | 0.6769 | 0.4699 | 0.3370 | 0.7495 | 0.5609 | 0.3599 | 0.05 | 0.3419 |
| | PIPR | 0.7844 | 0.5709 | 0.5526 | 0.4111 | 0.7720 | 0.7304 | 0.5355 | 0.07 | 0.4980 |
| | DeepFE-PPI | 0.7795 | 0.5743 | 0.5082 | 0.3726 | 0.7489 | 0.5415 | 0.4558 | 0.13 | 0.3965 |
| | GAT | 0.7996 | 0.6673 | 0.5379 | 0.3279 | 0.7836 | 0.5562 | 0.3441 | 0.04 | 0.3303 |
| 4 | HGNNPIP | 0.8504 | 0.6618 | 0.6046 | 0.4828 | 0.789 | 0.6138 | 0.5565 | 0.08 | **0.5120** |
| | DeepTrio | 0.8255 | 0.6391 | 0.4218 | 0.3299 | 0.7438 | 0.4930 | 0.3148 | 0.02 | 0.3085 |
| | PIPR | 0.8155 | 0.5305 | 0.5114 | 0.3982 | 0.7713 | 0.7140 | 0.4936 | 0.05 | 0.4689 |
| | DeepFE-PPI | 0.8180 | 0.5750 | 0.4319 | 0.3467 | 0.7507 | 0.4897 | 0.3458 | 0.10 | 0.3112 |
| | GAT | 0.8168 | 0.6517 | 0.3116 | 0.2597 | 0.78 | 0.4997 | 0.2048 | 0.05 | 0.1946 |
| 5 | HGNNPIP | 0.8807 | 0.6592 | 0.5938 | 0.4968 | 0.8155 | 0.6323 | 0.5401 | 0.05 | **0.5131** |
| | DeepTrio | 0.8482 | 0.6116 | 0.4207 | 0.3450 | 0.7532 | 0.4636 | 0.3206 | 0.03 | 0.3110 |
| | PIPR | 0.8360 | 0.5190 | 0.5047 | 0.4068 | 0.7869 | 0.7160 | 0.4912 | 0.03 | 0.4765 |
| | DeepFE-PPI | 0.8510 | 0.6232 | 0.3758 | 0.3410 | 0.7629 | 0.4657 | 0.2690 | 0.03 | 0.2609 |
| | GAT | 0.8397 | 0.68 | 0.1594 | 0.1845 | 0.7719 | 0.4526 | 0.0903 | 0.00 | 0.0903 |

**Table 5**. The predictions of five models on the testing set NTS2. All models are trained using the corrected datasets. The negative instances are generated by RanNS and the number is 2 folds of positive PPIs ($k = 2$).

| Dataset | Model | Subcellular localization | Random negative sampling |
|---|---|---|---|
| *S. cerevisiae* | HGNNPIP | 1 | 0.07 |
| | DeepTrio | 1 | 0.07 |
| | PIPR | 0.84 | 0.15 |
| | DeepFE-PPI | 0.96 | 0.17 |
| | GAT | 0.85 | 0.13 |
| *C. elegans* | HGNNPIP | 1 | 0.16 |
| | DeepTrio | 1 | 0.13 |
| | PIPR | 0.98 | 0.13 |
| | DeepFE-PPI | 1 | 0.16 |
| | GAT | 0.89 | 0.11 |
| *E. coli* | HGNNPIP | 0.96 | 0.08 |
| | DeepTrio | 0.98 | 0.02 |
| | PIPR | 0.97 | 0.12 |
| | DeepFE-PPI | 0.82 | 0.08 |
| | GAT | 0.90 | 0.04 |
| *D. melanogaster* | HGNNPIP | 1 | 0.17 |
| | DeepTrio | 1 | 0.13 |
| | PIPR | 1 | 0.15 |
| | DeepFE-PPI | 1 | 0.18 |
| | GAT | 0.95 | 0.23 |
| *Human* | HGNNPIP | 0.98 | 0.02 |
| | DeepTrio | 0.98 | 0.04 |
| | PIPR | 0.95 | 0.21 |
| | DeepFE-PPI | 0.97 | 0.13 |
| | GAT | 0.92 | 0.08 |
| *O. sativa* | HGNNPIP | 1 | 0.17 |
| | DeepTrio | 1 | 0.55 |
| | PIPR | 0.91 | 0.27 |
| | DeepFE-PPI | 0.98 | 0.55 |
| | GAT | 0.96 | 0.38 |

**Table 6**. The parameter optimization with dataset *S. cerevisiae*. The parameter $m$ denotes the length of the truncated subsequence in sequence encoding module. R_dim: the embedded dimensionality of each amino-acid residue; Dropout refers to disregard certain nodes at random during network training (range is from 0 to 1). DNN_outdim: the dimensionality of sequence feature. GAT_outdim: the dimensionality of topological feature. GAT_hiddim: the dimensionality of hidden layer in GAT; MLP_outdim: the number of neuros in the last layers of the MLP model.

| Round | $m$ | R_dim | Dropout | DNN_outdim | GAT_hiddim | GAT_outdim | MLP_outdim | Accuracy |
|---|---|---|---|---|---|---|---|---|
| 1 | 70 | 6 | 0.4 | 32 | 256 | 32 | 16 | 0.804 |
| 2 | 150 | 6 | 0.5 | 16 | 256 | 16 | 8 | 0.76 |
| 3 | 150 | 3 | 0.5 | 16 | 128 | 16 | 16 | 0.768 |
| 4 | 100 | 6 | 0.5 | 32 | 128 | 32 | 16 | 0.794 |
| 5 | 100 | 3 | 0.5 | 16 | 256 | 16 | 8 | 0.76 |
| **6** | **150** | **6** | **0.4** | **32** | **128** | **32** | **16** | **0.807** |
| 7 | 150 | 6 | 0.5 | 16 | 256 | 32 | 8 | 0.771 |
| 8 | 150 | 3 | 0.5 | 32 | 128 | 16 | 16 | 0.782 |
| 9 | 70 | 6 | 0.5 | 32 | 256 | 16 | 8 | 0.778 |
| 10 | 100 | 6 | 0.5 | 16 | 256 | 32 | 8 | 0.781 |
| 11 | 100 | 3 | 0.5 | 32 | 128 | 32 | 8 | 0.789 |
| 12 | 70 | 6 | 0.4 | 32 | 256 | 32 | 16 | 0.803 |
| 13 | 70 | 3 | 0.4 | 32 | 256 | 32 | 16 | 0.796 |
| 14 | 70 | 6 | 0.4 | 32 | 256 | 32 | 16 | 0.805 |
| 15 | 70 | 6 | 0.4 | 32 | 256 | 32 | 16 | 0.802 |

**Table 7**. Model output evaluation with network only, sequence only, and both. The fold of negative /positive is 2 ($k = 2$).

| Dataset | Model | Accuracy | Precision | Recall | Specificity | F1-score | MCC | AUROC | AP |
|---|---|---|---|---|---|---|---|---|---|
| *S. cerevisiae* | Both | **0.7964** | **0.7268** | 0.6246 | **0.8824** | **0.6716** | **0.5288** | **0.8260** | **0.7406** |
| | Sequence only | 0.7152 | 0.5678 | 0.6134 | 0.7664 | 0.5894 | 0.3450 | 0.7376 | 0.6030 |
| | Network only | 0.7282 | 0.5752 | **0.7160** | 0.7344 | 0.6374 | 0.4075 | 0.7910 | 0.6834 |
| *C. elegans* | Both | **0.8128** | **0.7244** | 0.7064 | **0.8654** | **0.7150** | **0.5758** | **0.8508** | **0.7780** |
| | Sequence only | 0.7648 | 0.6360 | 0.6890 | 0.8028 | 0.6612 | 0.4376 | 0.7964 | 0.7138 |
| | Network only | 0.6902 | 0.5242 | **0.7682** | 0.6506 | 0.6230 | 0.3880 | 0.7852 | 0.6776 |
| *E. coli* | Both | **0.8590** | **0.8348** | 0.7196 | **0.9288** | **0.7726** | **0.6752** | **0.8860** | **0.8492** |
| | Sequence only | 0.8338 | 0.7846 | 0.6914 | 0.9050 | 0.7348 | 0.5370 | 0.8078 | 0.7938 |
| | Network only | 0.8374 | 0.7704 | **0.7302** | 0.8910 | 0.7496 | 0.5600 | 0.8532 | 0.8256 |
| *D. melanogaster* | Both | **0.7860** | **0.6958** | 0.6279 | **0.8641** | **0.6601** | **0.5061** | **0.8199** | **0.7265** |
| | Sequence only | 0.7782 | 0.6780 | 0.6406 | 0.8468 | 0.6578 | 0.4338 | 0.8144 | 0.7214 |
| | Network only | 0.7060 | 0.5420 | **0.7702** | 0.6738 | 0.6358 | 0.4066 | 0.8034 | 0.7145 |
| *Human* | Both | **0.8398** | **0.7653** | 0.7522 | **0.8841** | **0.7583** | **0.6388** | **0.8910** | **0.8412** |
| | Sequence only | 0.7985 | 0.7114 | 0.6688 | 0.8640 | 0.6890 | 0.4744 | 0.8360 | 0.7620 |
| | Network only | 0.7628 | 0.6088 | **0.8086** | 0.7400 | 0.6944 | 0.4948 | 0.8514 | 0.7965 |
| *O. sativa* | Both | **0.9102** | **0.8542** | **0.8818** | **0.9248** | **0.8678** | **0.8004** | **0.9618** | **0.9392** |
| | Sequence only | 0.8222 | 0.7140 | 0.7792 | 0.8440 | 0.7452 | 0.5602 | 0.8928 | 0.8056 |
| | Network only | 0.8336 | 0.7084 | 0.8520 | 0.8248 | 0.7736 | 0.6154 | 0.9144 | 0.8618 |

**Table 8**. Performance Comparison on six reconstructed datasets. The fold of negative /positive is 2 ($k = 2$).

| Dataset | Model | Accuracy | Precision | Recall | Specificity | F1-score | MCC | AUROC | AP |
|---|---|---|---|---|---|---|---|---|---|
| *S. cerevisiae* | HGNNPIP | **0.7964** | **0.7268** | **0.6246** | 0.8824 | **0.6716** | **0.5288** | **0.826** | **0.7406** |
| | DeepTrio | 0.7582 | 0.6944 | 0.4688 | **0.8982** | 0.5561 | 0.3401 | 0.7612 | 0.6417 |
| | PIPR | 0.7315 | 0.5981 | 0.5853 | 0.7958 | 0.5916 | 0.3917 | 0.7601 | 0.730 |
| | DeepFE-PPI | 0.7303 | 0.6057 | 0.5472 | 0.8219 | 0.5748 | 0.3792 | 0.7471 | 0.6103 |
| | GAT | 0.7178 | 0.5724 | 0.6068 | 0.7732 | 0.5890 | 0.3748 | 0.7342 | 0.6102 |
| *C. elegans* | HGNNPIP | **0.8128** | 0.7244 | **0.7064** | 0.8654 | **0.7150** | **0.5758** | **0.8508** | **0.778** |
| | DeepTrio | 0.8038 | **0.7845** | 0.6019 | **0.9105** | 0.6781 | 0.4544 | 0.8301 | 0.7584 |
| | PIPR | 0.7600 | 0.6413 | 0.6230 | 0.8163 | 0.6319 | 0.4541 | 0.8081 | 0.786 |
| | DeepFE-PPI | 0.7451 | 0.6200 | 0.6102 | 0.8125 | 0.6140 | 0.4248 | 0.7821 | 0.6635 |
| | GAT | 0.7252 | 0.5990 | 0.5364 | 0.8200 | 0.5658 | 0.3674 | 0.731 | 0.6198 |
| *E. coli* | HGNNPIP | **0.859** | 0.8348 | **0.7196** | 0.9288 | **0.7726** | **0.6752** | **0.886** | 0.8492 |
| | DeepTrio | 0.8468 | **0.8564** | 0.6198 | **0.9517** | 0.7190 | 0.5322 | 0.8376 | 0.8005 |
| | PIPR | 0.8479 | 0.7834 | 0.7575 | 0.8791 | 0.7701 | 0.6568 | 0.8879 | **0.8729** |
| | DeepFE-PPI | 0.8191 | 0.7653 | 0.6611 | 0.8981 | 0.7089 | 0.5823 | 0.8185 | 0.7733 |
| | GAT | 0.8072 | 0.7160 | 0.6998 | 0.8608 | 0.7076 | 0.564 | 0.8108 | 0.7902 |
| *D. melanogaster* | HGNNPIP | **0.7860** | 0.6958 | **0.6279** | 0.8641 | **0.6601** | **0.5061** | 0.8199 | 0.7265 |
| | DeepTrio | 0.7850 | **0.7127** | 0.6084 | **0.8746** | 0.6553 | 0.4295 | **0.8336** | 0.7373 |
| | PIPR | 0.7403 | 0.6132 | 0.6041 | 0.8027 | 0.6086 | 0.4144 | 0.7759 | **0.7507** |
| | DeepFE-PPI | 0.7539 | 0.6405 | 0.5978 | 0.8320 | 0.6181 | 0.4377 | 0.7876 | 0.6691 |
| | GAT | 0.7396 | 0.5987 | 0.6626 | 0.7784 | 0.6294 | 0.4310 | 0.7788 | 0.6770 |
| *Human* | HGNNPIP | **0.8398** | **0.7653** | **0.7522** | 0.8841 | **0.7583** | **0.6388** | **0.8910** | **0.8412** |
| | DeepTrio | 0.8141 | 0.7531 | 0.6495 | **0.8950** | 0.6968 | 0.4894 | 0.8587 | 0.7816 |
| | PIPR | 0.7825 | 0.6784 | 0.6677 | 0.8338 | 0.6729 | 0.5102 | 0.8304 | 0.8113 |
| | DeepFE-PPI | 0.7995 | 0.7148 | 0.6635 | 0.8674 | 0.6880 | 0.5415 | 0.8431 | 0.7608 |
| | GAT | 0.7856 | 0.6606 | 0.7346 | 0.8112 | 0.6952 | 0.5326 | 0.8376 | 0.7658 |
| *O. sativa* | HGNNPIP | **0.9102** | **0.8542** | **0.8818** | **0.9248** | **0.8678** | **0.8004** | **0.9618** | **0.9392** |
| | DeepTrio | 0.8467 | 0.8110 | 0.7025 | 0.9181 | 0.7520 | 0.5635 | 0.9163 | 0.8552 |
| | PIPR | 0.8726 | 0.8054 | 0.8125 | 0.9064 | 0.8089 | 0.7134 | 0.9290 | 0.9126 |
| | DeepFE-PPI | 0.8426 | 0.7653 | 0.7627 | 0.8825 | 0.7636 | 0.6460 | 0.9023 | 0.8273 |
| | GAT | 0.8284 | 0.7238 | 0.7850 | 0.8504 | 0.7532 | 0.6232 | 0.8884 | 0.82 |

**Table 9.** The interacting partners of OsEDS1 predicted by HGNNPIP model.

| Protein name | Interacting partners | Score |
|---|---|---|
| OsEDS1 | OsVQ13 | 0.988104 |
| OsEDS1 | OsWRKY45 | 0.952316 |
| OsEDS1 | OsHOX27 | 0.884054 |
| OsEDS1 | OsCUL1-3 | 0.882917 |
| OsEDS1 | OsRDBF | 0.879466 |
| OsEDS1 | Os09g0249000 | 0.869282 |
| OsEDS1 | OsDRB1a | 0.861887 |
| OsEDS1 | Os09g0465400 | 0.800382 |
| OsEDS1 | OsFer2 | 0.770617 |
| OsEDS1 | OsbZIP77 | 0.760558 |
| OsEDS1 | OsWRKY72 | 0.660121 |
| OsEDS1 | OsSCAR3 | 0.631592 |
| OsEDS1 | OsBTBZ4 | 0.630277 |
| OsEDS1 | OsHOX11 | 0.536224 |
| OsEDS1 | OsPCF8 | 0.523366 |

**Table 10.** The interacting partners of SCRE4 predicted by HGNNPIP model.

| Protein name | Interacting partners | Score |
|---|---|---|
| **SCRE4** | **OsRLCK33** | **0.634370** |
| **SCRE4** | **OsCML11** | **0.530550** |
| SCRE4 | OsCDPK6 | 0.186775 |
| SCRE4 | Os08g0558100 | 0.025386 |
| SCRE4 | OsCaM61 | 0.009076 |
| SCRE4 | OsCaM1 | 0.005669 |
| SCRE4 | OsCML2 | 0.005521 |
| SCRE4 | OsCML3 | 0.003829 |
| SCRE4 | OsCML24 | 0.001423 |
| SCRE4 | OsCDPK1 | 0.000465 |
| SCRE4 | OsCML32 | 0.000135 |
| SCRE4 | OsCML16 | 0.000127 |
| SCRE4 | OsCML21 | 2.66E-05 |
| SCRE4 | OsCDPK2 | 2.31E-05 |
| SCRE4 | OsWRKY53 | 1.91E-05 |
| SCRE4 | OsCPK22 | 1.79E-05 |
| SCRE4 | OsWRKY35 | 1.28E-05 |
| SCRE4 | OsCPK25 | 5.50E-06 |
| SCRE4 | OsCPK26 | 5.50E-06 |
| SCRE4 | OsCPK21 | 3.03E-06 |
| SCRE4 | OsCPK29 | 1.52E-06 |
| SCRE4 | OsRboh9 | 7.23E-07 |
| SCRE4 | OsCML27 | 6.18E-07 |
| SCRE4 | OsRbohB | 2.19E-07 |

**Table 11**HGNNPIP's performances on dataset *S. cerevisiae* with different classifiers (5-fold Cross-Validation).

| Model | Accuracy | Precision | Recall | Specificity | F1-score | MCC | AUROC | AP |
|---|---|---|---|---|---|---|---|---|
| MLP | **0.7964** | 0.7268 | **0.6246** | 0.8824 | **0.6716** | **0.5288** | **0.826** | **0.7406** |
| LR | 0.7130 | 0.5676 | 0.5828 | 0.7782 | 0.5750 | 0.3584 | 0.730 | 0.5892 |
| RF | 0.7900 | **0.8424** | 0.4496 | **0.9582** | 0.5862 | 0.5043 | 0.8252 | 0.7382 |
| SVM | 0.7800 | 0.6956 | 0.6080 | 0.8662 | 0.6488 | 0.4903 | 0.801 | 0.7098 |
