## Supplementary materials for "HGNNPIP: A Hybrid Graph Neural Network framework for Protein-protein Interaction Prediction"

### Supplementary Figures

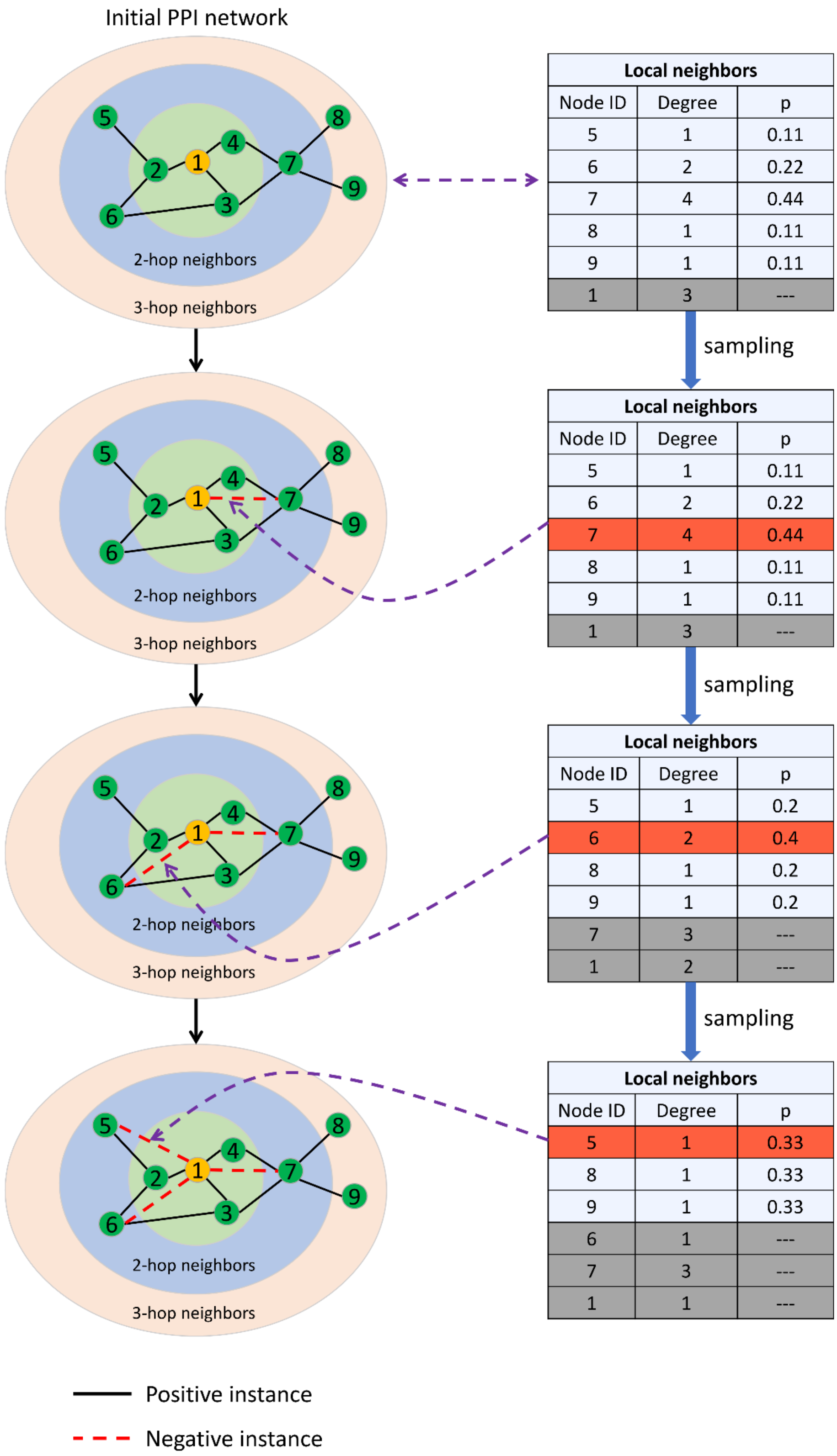


**Supplementary Fig. 1** The process of popularity-based negative sampling strategy.

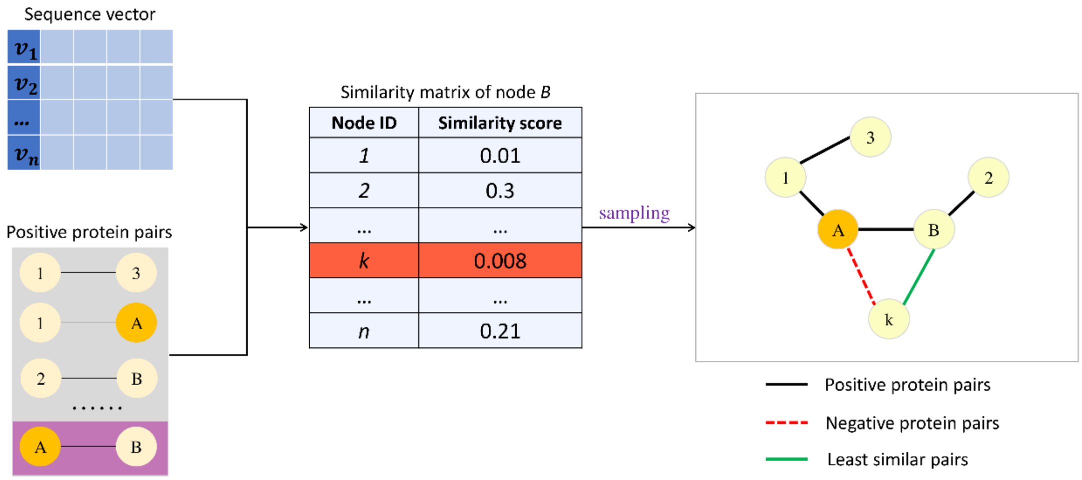


**Supplementary Fig. 2** The diagram of similarity-biased negative sampling strategy.

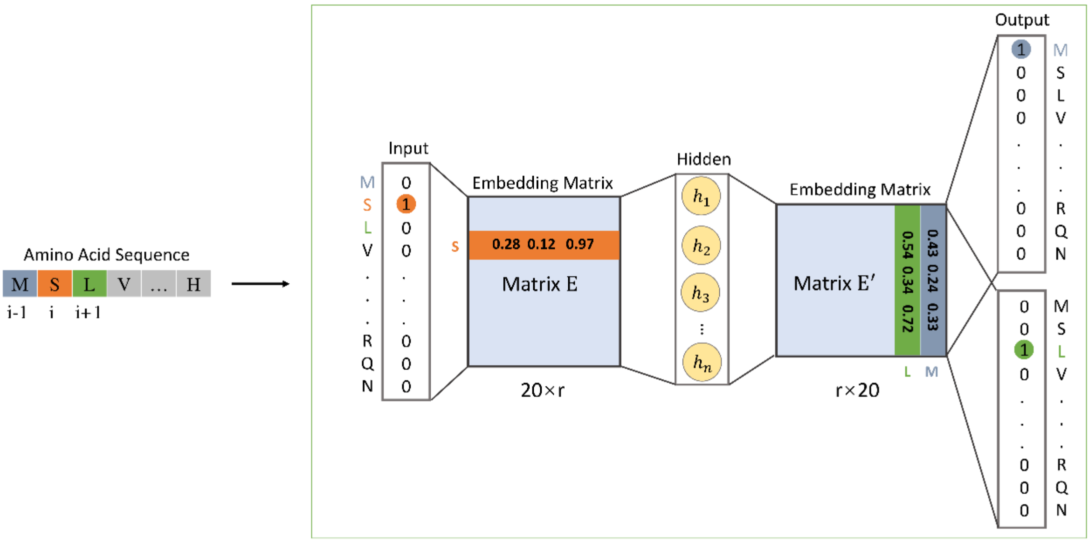


**Supplementary Fig. S3** The process of training the Word2vec model for amino acid residues.

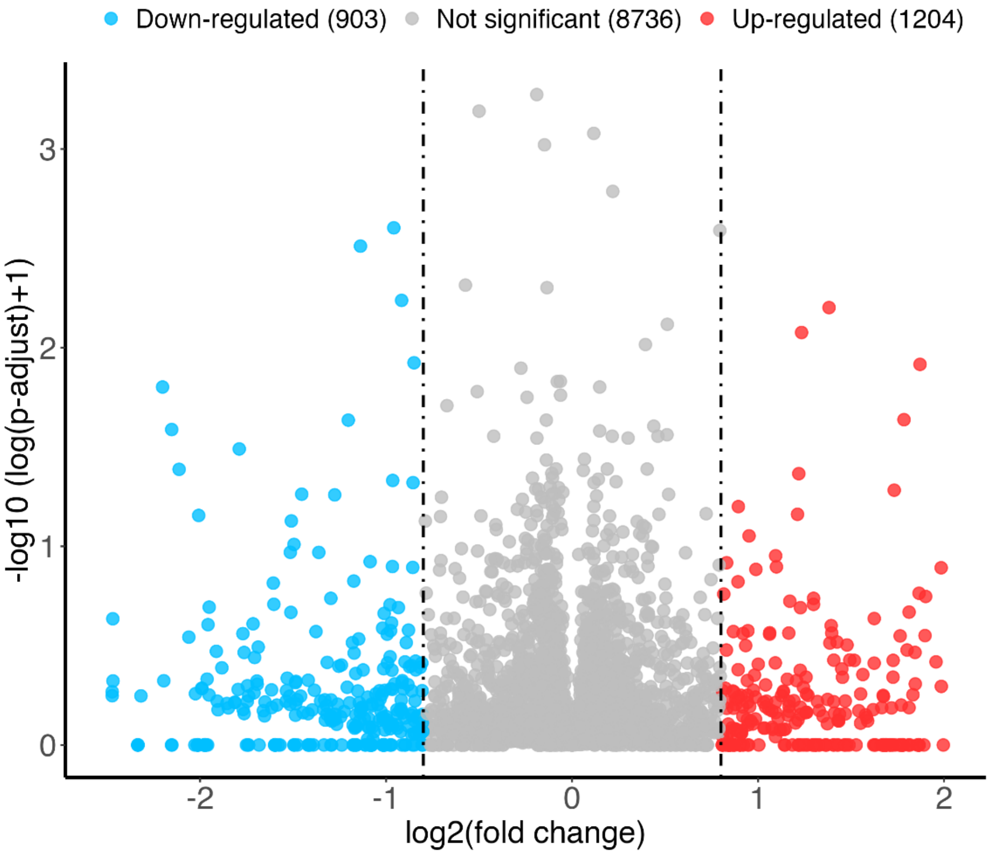


**Supplementary Fig. 4**. Volcano plots of DEGs analysis. The fold changes (FC) of (OsEDS1-mutant vs. control) are evaluated. Differential expressed genes are determined if the genes follow the criterion: |logFC| > 0.8 and adjusted P-value < 0.05.

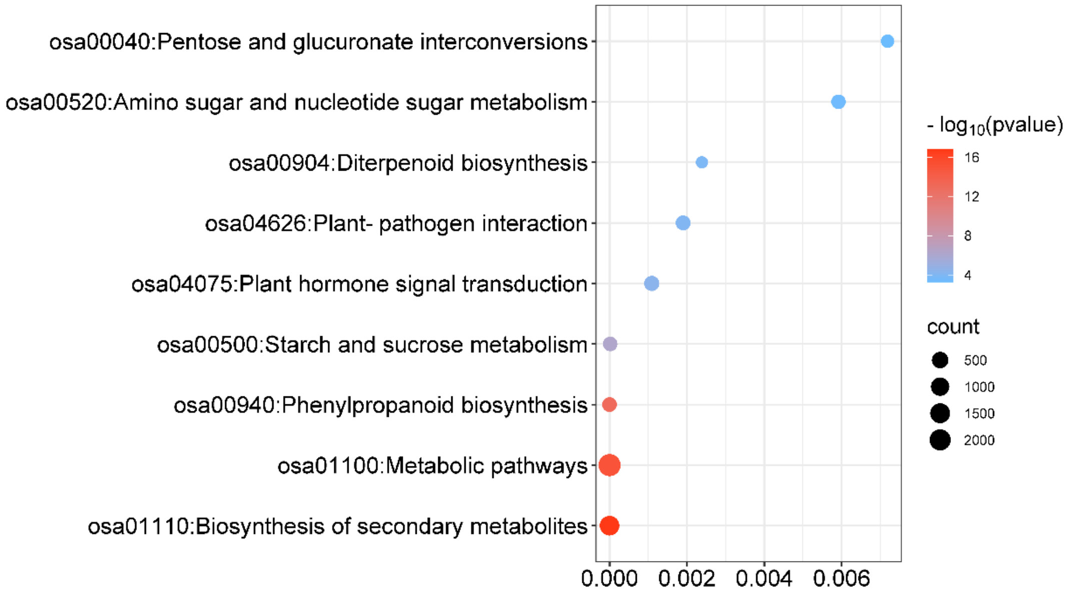


**Supplementary Fig. 5** Pathway enrichment analysis for the transcriptome data of rice with false smut.

### Supplementary Tables

**Supplementary Table 1.** Primer sets used in this study.

| Accession number | Primer name | Primer sequence |
|---|---|---|
| Os06g0255200 | Os06g0255200-qF | ATGGATGCGTTGCGATCATC |
| | Os06g0255200-qR | ATGCGCACTACTTTGGGATG |
| Os10g0561800 | OsHOX1-qF | GGACACCTTCAAAGAGCACAAC |
| | OsHOX1-qR | TCTGCTTCAGCTTCGTCCTC |
| Os05g0186900 | OsIAA16-qF | ATGGCAGGTTTGGAGAGGATG |
| | OsIAA16-qR | GCAGGAAACTTTGAGGGCAAG |
| Os12g0616400 | OsPCF8-qF | AAGCCATTGCAGCAACAGC |
| | OsPCF8-qR | ATCACCTCGAACGACATGGG |
| Os01g0656400 | OsWRKY15-qF | TTGCCTCCAAGACGAACTCC |
| | OsWRKY15-qR | AGGTGTCCAAGAAATCGGAGTC |
| Os03g0676400 | OsVQ13-qF | CGGACATGTTCGACTACGC |
| | OsVQ13-qR | ACAGTAAAGCACCATGTCCATG |
| Os02g0149600 | Os02g0149600-qF | TGCAGCAACAATCACCAGAG |
| | Os02g0149600-qR | AGTGGGCATTTCAACGAAGG |
| Os11g0490900 | OsWRKY72-qF | AGAAGGCCGTCAAGAACAAC |
| | OsWRKY72-qR | TTCTTCACGTTGCACCCTTG |
| Os02g0770800 | OsNIA1-qF | TGAAATGGCTCAAGCGCATC |
| | OsNIA1-qR | ATCATGTACTCCGGCTTGTACC |
| Os02g0252400 | RPBF-qF | AGCTGTGATGACGAAGGACAC |
| | RPBF-qR | TGTTCCAGCCGTAGAAGTAGTC |
| Os09g0249000 | Os09g0249000-qF | ATGTACGCCTTCGACCTCAAG |
| | Os09g0249000-qR | TTCAGCTCCTTGTGCTCCTG |
| Os01g0883100 | OsMADS2-qF | AAACTCTCTGCAGCCCAAAG |
| | OsMADS2-qR | AGCAGCTTGTTCTCGTCTTC |
| Os05g0322900 | OsWRKY45-qF | TTCCTTGTTGATGTGTCGTCTCA |
| | OsWRKY45-qR | CCCCCAGCTCATAATCAAGAAC |
| Os03g0718100 | OsActin-F | TGTATGCCAGTGGTCGTACCA |
| | OsActin-R | CCAGCAAGGTCGAGACGAA |
